# A novel domain assembly routine for creating full-length models of membrane proteins from known domain structures

**DOI:** 10.1101/209213

**Authors:** Julia Koehler Leman, Richard Bonneau

## Abstract

Membrane proteins composed of soluble and membrane domains are often studied one domain at a time. However, to understand the biological function of entire protein systems and their interactions with each other and drugs, knowledge of full-length structures or models is required. Although few computational methods exist that could potentially be used to model full-length constructs of membrane proteins, none of these methods are perfectly suited for the problem at hand. Existing methods either require an interface or knowledge of the relative orientations of the domains, are not designed for domain assembly, and none of them are developed for membrane proteins. Here we describe the first domain assembly protocol specifically designed for membrane proteins that assembles intra- and extracellular soluble domains and the transmembrane domain into models of the full-length membrane protein. Our protocol does not require an interface between the domains and samples possible domain orientations based on backbone dihedrals in the flexible linker regions, created via fragment insertion, while keeping the transmembrane domain fixed in the membrane. Our method, mp_domain_assembly, implemented in RosettaMP samples domain orientations close to the native structure and is best used in conjunction with experimental data to reduce the conformational search space.

## Introduction

It is well established that membrane proteins are very difficult to study experimentally, which is why many laboratories cleave transmembrane domains (TMD) from their protein constructs and determine structures of the soluble domains in isolation. Structures of the TMD are elucidated separately, mostly by groups specializing in membrane proteins. However, to gain insights into the biological workings of complex and intricate protein systems, models of full-length constructs are needed, which can be used to generate hypotheses about protein function and their interactions with other proteins or small molecules. While crystallography and cryo-EM have advanced to the point where larger constructs spanning soluble and TMDs are determined, systems with flexible linkers constitute a major challenge for structure determination.

Further, proteins with single transmembrane spans are difficult to study by crystallography, yet constitute about half of the membrane proteins in the genome^1,2^. Prime examples are receptor tyrosine kinases (such as the insulin receptor, epidermal growth factor receptor (EGFR), fibroblast growth factor receptor (FGFR)), adhesion molecules such as integrins, β-subunits of voltage-gated potassium channels, sodium channels and calcium channels, and others. For example, one of the long-standing questions in structural biology is how receptor tyrosine kinases signal across the membrane bilayer to lead to downstream signaling effects^3^. Availability of a full-length model would accelerate progress in this field and facilitate drug development for a variety of diseases including many types of cancer^4,5^.

Few domain assembly protocols have been developed over the years, and they are mainly docking protocols that find the optimal relative orientation of interacting domains by sampling flexible linkers. Wollacott et al. developed a method where the flexible linker was sampled via fragment insertion in low-resolution mode^6^. Models with RMSDs < 3 Å to the reference structure in the benchmark set were subject to a high-resolution step where optimal solutions were found by examining the interface energy between the two domains. RosettaRemodel^7^ was recently developed for a variety of modeling problems, including domain assembly and loop modeling. It was used for domain insertion of a specific protein in between two linkers, which substantially reduced the conformational search space. However, RosettaRemodel is most often used for loop modeling^8,9^ and design applications^10^. Godzik and co-workers recently developed AIDA^11^ (*ab initio* domain assembly) where the individual domains are first modeled with Modeller and then assembled. Model quality is evaluated based on the interface energy between the two domains. It seems that most tested proteins had large interfaces, and proteins without contacts between the individual domains were removed from the benchmark set. The authors find that the success rate to find native-like models drops drastically for longer linkers (~50% for 6-10 residue linkers and ~25% for 11-20 residue linkers)^11^.

Loop modeling protocols have been used for assembling membrane protein domains in a manual fashion^12,13^. However, both kinematic loop closure^8,14^ as well as cyclic coordinate descent^15,16^ can only be used to model loops between fixed endpoints. Rosetta’s FloppyTail protocol^17^ is able to model flexible termini using Small and ShearMovers and 3-residue fragments, yet is not designed to assemble domains in an automated fashion. FloppyTail was used to build models of the flexible tail of Cdc34 and the resulting models were filtered based on the interaction energy between the terminus and its interaction partner^17^.

While the methods described above are undoubtedly useful, none of them really fit the problem at hand: (1) the existing domain assembly protocols assume a sufficiently large interface between the individual domains and use the interface energy for scoring or to drive the domain assembly. In our case, the different domains often have no or only a very small interface. (2) Loop modeling or RosettaRemodel methods model flexible loops between fixed endpoints in space. This requires knowing the relative orientations of the domains, ignoring the underlying problem of flexible linkers. Our questions are: “*what are possible conformations of the flexible linker regions?*” and “*what relative orientations of the domains can be sampled under these constraints?*” (3) FloppyTail cannot assemble domains in an automated fashion. (4) None of these methods are designed for membrane proteins, requiring that the TMD remains fixed in the membrane, while the N- and C-terminal domains can move around flexibly. In the Rosetta software, we accomplish this via a specific setup of the FoldTree^16^, which assembles the domains from the center of the membrane as the starting point (see below) and updates coordinates in a tree-like fashion by perturbing backbone dihedrals towards the N- and C-termini in dihedral angle space. Further, we wanted to develop a protocol that is easily usable by non-expert modelers.

Here, we describe a Monte-Carlo domain assembly protocol that can be used to create full-length models of membrane proteins from structures or models of its domains, whether they are in solution or in the membrane. This method is the first designed specifically for membrane proteins. Our protocol samples linker dihedral angles from fragments of known protein structures, thereby sampling linker conformations that have been seen in nature. This application is implemented in the Rosetta software suite^18^, is based on the recently implemented RosettaMP^19^ framework, and we provide command lines and a protocol capture to ensure usability for non-expert modelers.

## Development of the method

Our protocol *mp_domain_assembly* requires the full-length sequence in fasta format, 9- and 3-residue fragments that can be created with the Robetta server^20^ or the fragment picking pipeline ^21^, and the structures or models of the domains of interest. The user further has to specify on the command line which of the structures is the TMD. The structures are read into Rosetta and the sequences from the structures’ ATOM lines are aligned with the sequence from the fasta file to identify the linker residues (FIG 1A).

**Fig. 1:**
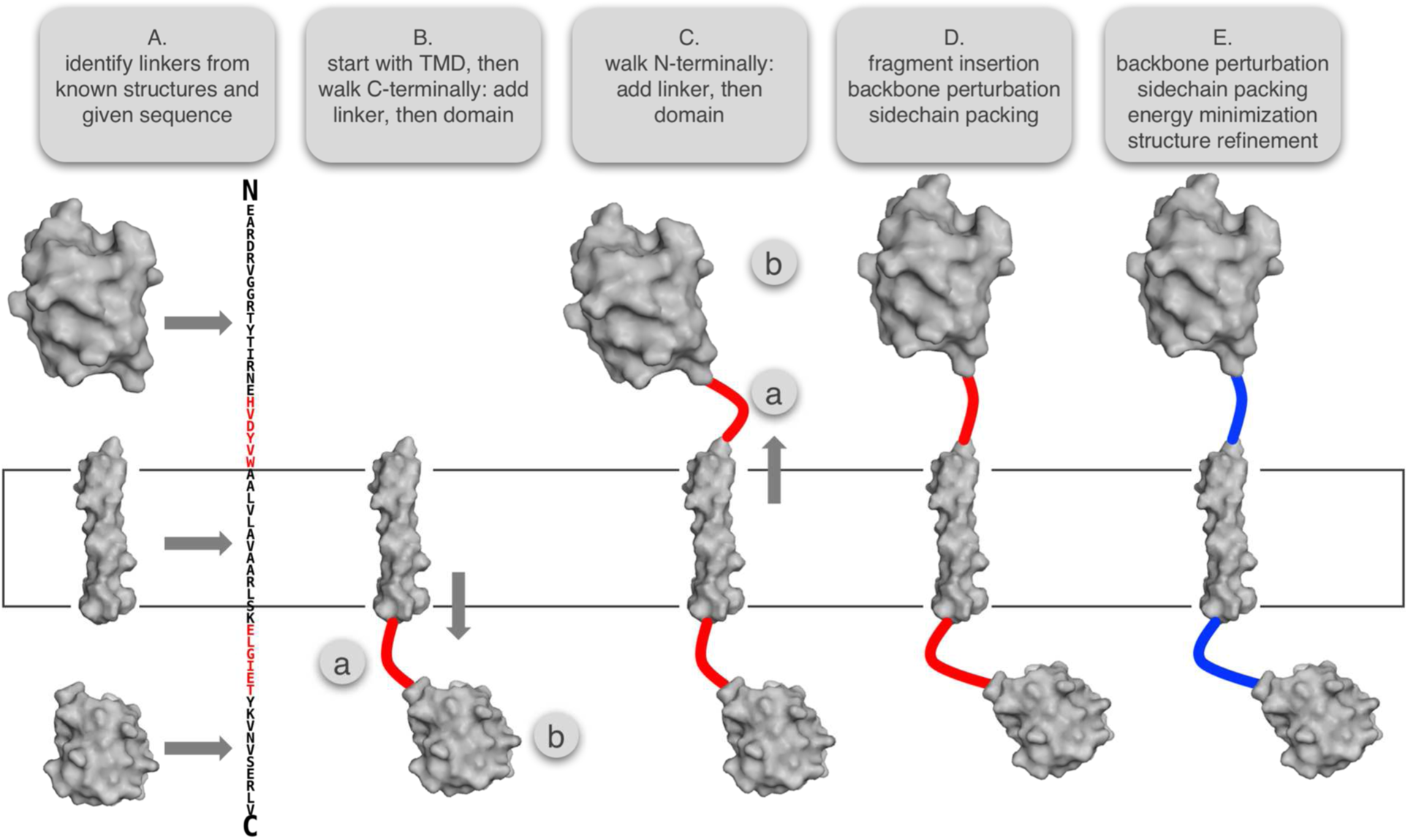
Steps for the domain assembly protocol for membrane proteins. Required inputs are the structures or models of the individual domains, the sequence of the full-length construct and fragments (for instance generated with the Robetta server). Domain assembly requires that the transmembrane domain (TMD) is already embedded in the membrane (see protocol capture). Linker residues and the C-terminal domains are then added one-by-one. Walking N-terminally, both linker and domain are then attached. Linker conformations are then sampled via fragment insertion and dihedral angle perturbations, while optimizing sidechain rotamers. Final steps include dihedral angle perturbations, rotamer optimization and energy minimization, before an optional high-resolution refinement step of the full-length construct.

Domain assembly then starts from the TMD, which at this point is already embedded in the membrane bilayer (see prerequisites). Moving along the sequence in C-terminal direction, linker residues are added to the pose one-by-one with temporarily random torsion angles (FIG 1B). When a residue is found that is part of another domain, two dummy residues are added to the N-terminal end of this domain, these two residues are superimposed with the ones in the linker, and the dummy residues are then removed. A continuous full-length pose is thus built up in an iterative fashion: linker, domain, linker, domain, etc. After all domains are added from the TMD to the C-terminus, linker residues and domains are added moving backwards along the sequence (from the TMD to the N-terminus – FIG 1C).

The full-length pose is then used to create a *SpanningTopology* object which defines the transmembrane spans, and is required to add the virtual membrane residue via the *AddMembraneMover* in the RosettaMP framework^19^. The spanfile is written out during that step. During the setup of the membrane representation, we create a FoldTree^16^ object that has its root at the residue closest to the transmembrane center-of-mass. This ensures that the TMD remains fixed in the membrane while all N- and C-terminal domains can move around as rigid bodies, depending on the flexibility of the linker residues. The scorefunction we use is *mpframework_smooth_fa_2012* until an improved scorefunction has been developed.

The pose, while full-length at this point, has clashes because none of the dihedral angles in the linker residues have been optimized. We then create a MoveMap^22^ which allows only the linker residues +/-1 to move. In *n* cycles (below), we insert 9-residue fragments, then 3-residue fragments, apply a SmallMover (with maximal torsion perturbation of 180°) followed by sidechain optimization of the linker residues, and apply a ShearMover (with maximal torsion perturbation of 180°) followed by sidechain optimization of the linker residues; each step is evaluated by the Metropolis criterion (FIG 1D). The number of cycles *n* was empirically determined and seems to have an optimal value for (*n* = 8 · <total number of linker residues>), considering the fraction of accepted steps for fragment insertion and the runtime. More sampling did not result in better linker conformations on our benchmark, yet increased runtimes. The final steps in this protocol include another SmallMover and ShearMover for backbone perturbations, sidechain packing of the linker residues, and energy minimization (lbfgs_armijo_atol with a tolerance of 0.01 in a maximum of 2000 steps), each with evaluation of the Metropolis criterion in between (FIG 1E). The last step is an optional high-resolution refinement run of the entire construct to remove clashes and improve local geometry.

## Validation of the method

To test our method, we collected five available structures from the bitopic protein superfamilies in the OPM database^23^ that have a single TMD and a soluble domain. Most bitopic membrane proteins with determined structures are NMR structures of the TMD only, reinforcing our point of very few available structures. Our examples (Table 1) are the sigma intracellular receptor (PDBIDs 5hk1 and 5hk2, they have different linker conformations and respective domain orientations), the monoamine oxidase A (PDBID 2z5x), the penicillin-binding protein 1B (PDBID 5hlb), and the β1 subunit of the electric eel Nav1.4 sodium channel (PDBID 5xsy).

**Table 1:**
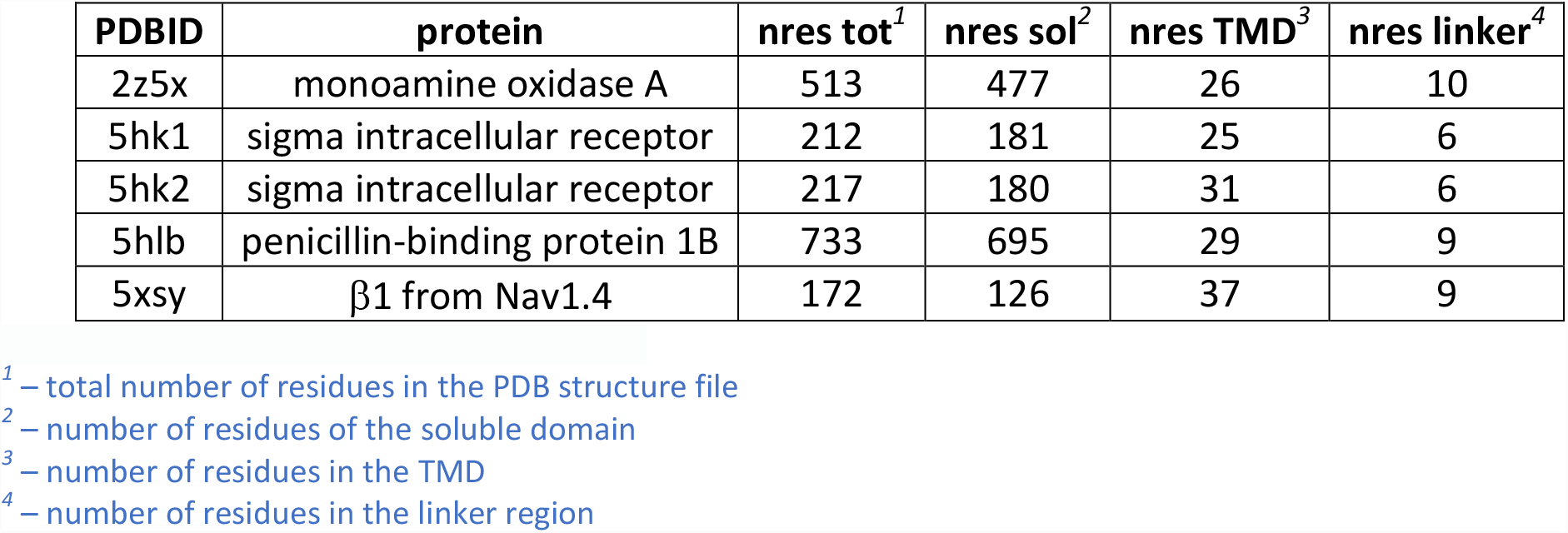
Examples in our benchmark dataset

Structures were downloaded from the PDBTM database^24^ and their membrane embedding was further optimized in Rosetta. The PDB files were divided into the TMD and the soluble domain by removing the linker residues. The number of linker residues ranged from 6 to 10 residues. Fragments were generated with Robetta^20^ for which homologues were excluded. We created 100,000 models for each protein, keeping the TMD fixed in the membrane while the soluble domains could move around flexibly depending on the backbone dihedral angles of the linker residues. Since the models are superimposed on the TMDs (which are single transmembrane span helices in this dataset) and the soluble domains contain many more residues, the computed RMSDs can become large due to the lever-arm effect.

Rosetta score vs. Cα-RMSD plots are shown in FIG 2 where the models in blue are the refined native structure. We find that in all test cases native domain orientations were recapitulated by our protocol. In FIG 1 we show the lowest RMSD models generated with this protocol. The lowest RMSD models are at 5.5 Å for 2z5x (FIG 2A), 2.8 Å for 5hk1 (FIG 2B) 3.5 Å for 5hlb (FIG 2C), and 5.1 Å for 5xsy (FIG 2D). While improvements to the membrane scorefunction might increase the ability to select native-like conformations, we find these results extremely encouraging. Further, model selection can be drastically improved with the availability of experimental data.

**Fig. 2:**
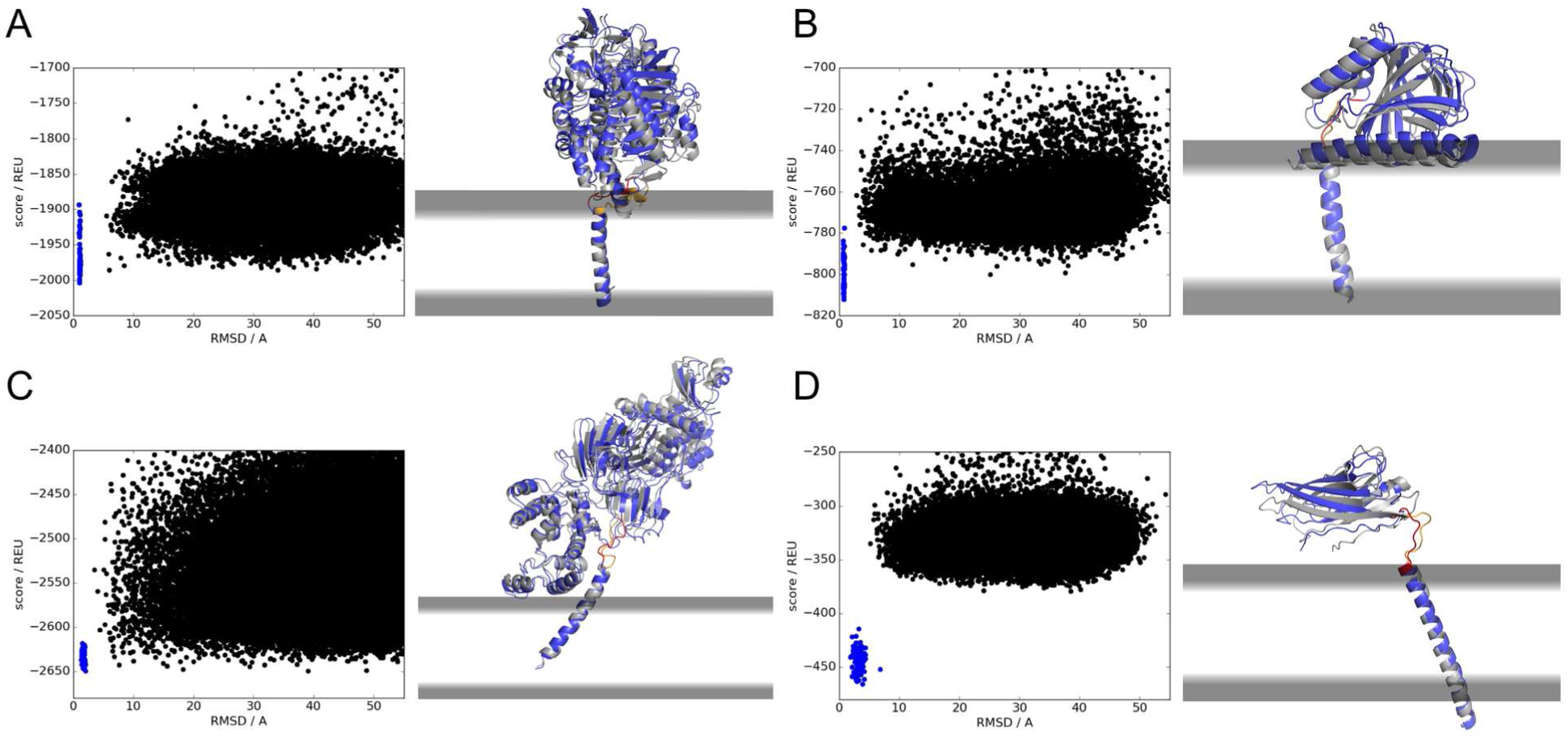
Examples used for benchmarking our method. Score vs Cα-RMSD plots are shown on the left, which contains the refined native structures as blue dots. The native structures in gray are superimposed with top RMSD models for (A) 2z5x with RMSD = 5.5 Å, (B) 5hk1 with RMSD = 2.8 Å, (C) 5hlb with RMSD = 3.5 Å, (D) 5xsy with RMSD = 5.1 Å. Native linkers are shown in yellow and modeled linkers are shown in red. The score vs RMSD plot and lowest RMSD model for 5hk2 are shown in the Supplement.

FIG 3 shows the runtimes for our protocol depending on the sequence length of the full-length construct. Runtimes depend on (1) the number of fragment insertions and therefore on the total number of residues in the linkers, (2) the number of residues in the soluble domains for which the coordinates need to be updated after the fragment insertion, and (3) the number of residues in the full-length construct for which minimization and scoring takes place. For the tested constructs runtimes remain on average below 32 minutes on a 2.7 GHz Intel Xeon E5 processor (FIG 3, black line). The most time-consuming step in this protocol is the optional high-resolution refinement step at the end of the protocol. Without refinement, full-length models can be built in under 250 seconds (~4 mins) even for the largest of the tested cases (FIG 3, gray line).

**Fig. 3:**
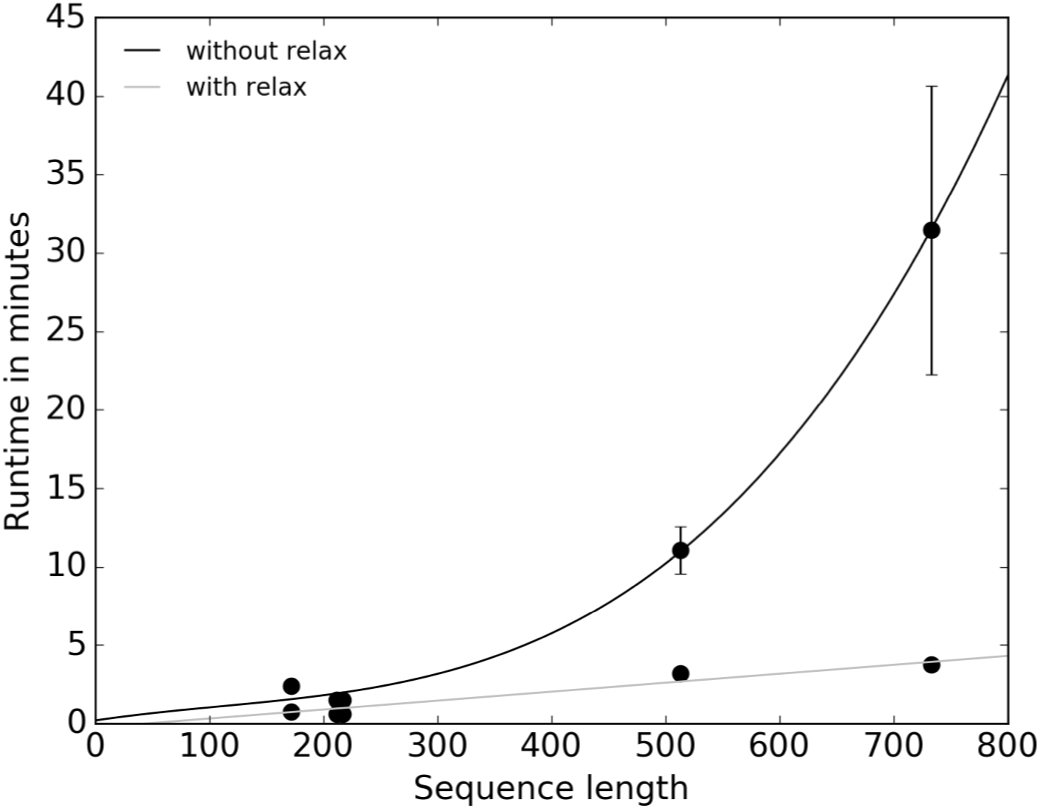
Runtimes for mp_domain_assembly depending on the sequence length of the assembled construct. Runtimes are shown with (black line) and without (gray line) the final refinement step in the protocol. Fitting parameters for the linear fit are: y = 0.36x-26.8. Our method is fast enough to assemble large to very large membrane proteins.

We used our algorithm to create full-length models of the fibroblast growth factor receptor 3 (FGFR3) monomer for which a structure does currently not exist. To accomplish this, we used the PDBID 4k33 for the tyrosine kinase domain. After cleaning and renumbering the PDB file, we mutated three residues back to their original wild-type residues (A24G, S124C, E188K where the latter is wild-type) and modeled a missing loop between residues 124 and 129 via KIC loop modeling^25^, generating 1000 models. We then refined each domain (PDBIDs 1ry7 for the Ig domain, 2lzl for the TMD, 4k33 for the tyrosine kinase domain) with our *range_relax* algorithm, generating 100 models for each. In each step, the top-scoring model was used for the next iteration. Refined models where then used for the domain assembly protocol as described in the protocol capture (see Supplement). The top 10 and top 1 scoring models are shown in FIG 4. The modeled linker lengths of FGFR3 were 9 and 59 residues, respectively. From the top 10 models, it is clear that experimental data is required to narrow down possible linker conformations or gain insights into how this protein functions biologically with such long disordered domains.

**Fig. 4:**
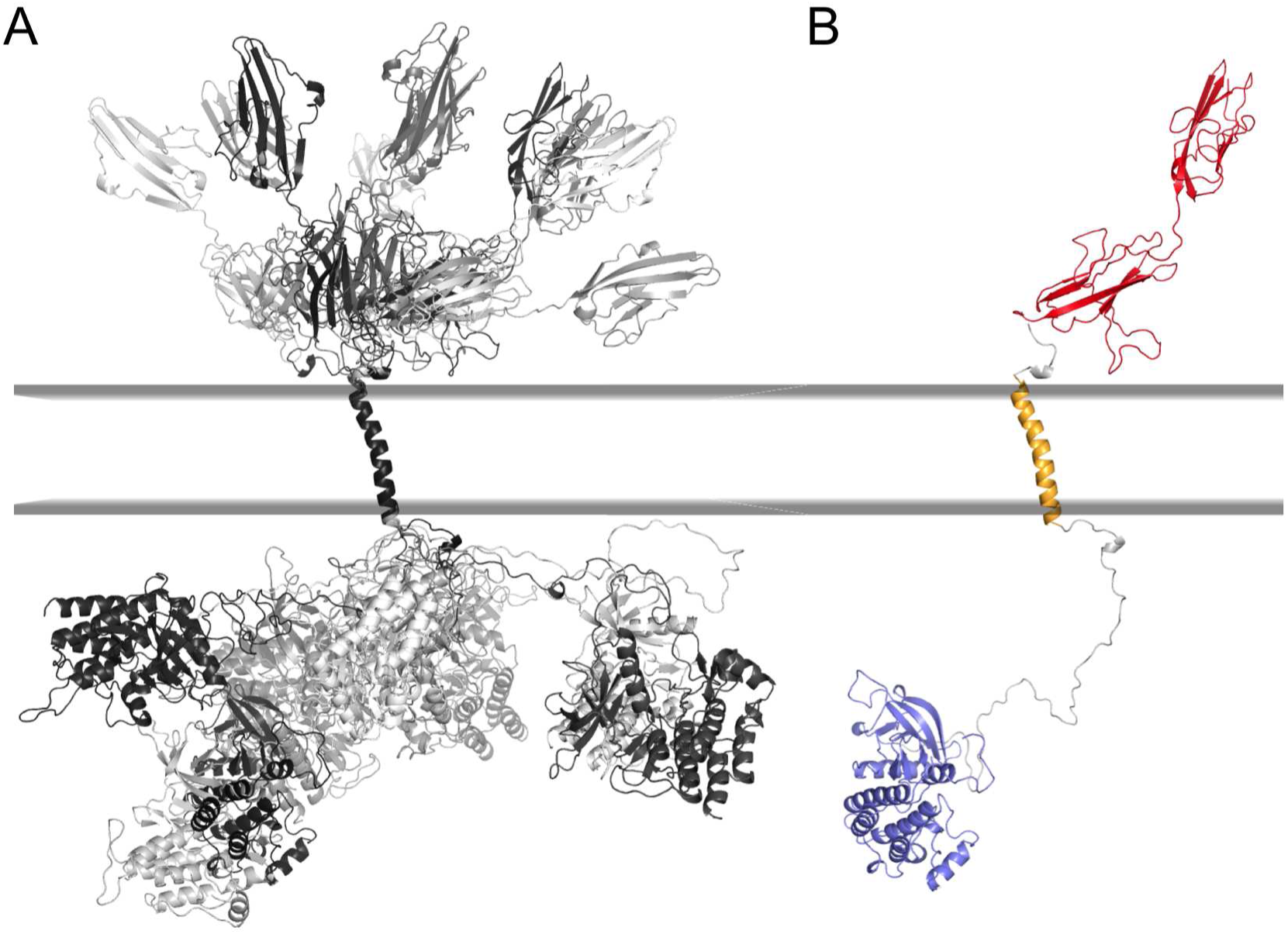
(A) Top 10 scoring models of 10,000 generated models of the fibroblast growth factor receptor 3 (FGFR3) monomer in different shades of gray. The models were generated from the domain structures of PDBIDs 1ry7, 2lzl, 4k33. The runtime to assemble the 606 residues was on average 276 +/-14 s without refinement. (B) Top scoring model of the FGFR3 monomer with the Ig domain in red, the TMD in yellow, the tyrosine kinase domain in blue, and the linkers in gray.

## Conclusion

Here we present the first domain assembly protocol specifically designed for membrane proteins. Modeling starts with the TMD remaining fixed in the membrane while linkers and soluble domains are attached in the C-terminal and then N-terminal directions to build up the full-length structure. Possible linker conformations are sampled via fragment insertion, dihedral angle perturbations and energy minimization. For five examples tested, our protocol is able to recapitulate domain orientations close to the native structure. Moreover, our method creates full-length models for proteins up to ~730 residues in, on average, under 4 minutes without refinement and in under 32 minutes with high-resolution refinement, enabling model building for large to very large proteins in a reasonable timeframe. We provide a protocol capture for usability by non-expert modelers and recommend using our method together with experimental data to reduce the conformational search space. The presented domain assembly method for membrane proteins would be particularly useful in conjunction with other hybrid structural biology approaches.

## Acknowledgements

The authors thank the Flatiron Institute at the Simons Foundation for funding and members of the RosettaCommons for helpful discussions.

## Supplementary Information

Supplementary information is available in the online version of this article. The file contains the protocol capture with the Rosetta command lines for the *mp_domain_assembly* protocol.

---

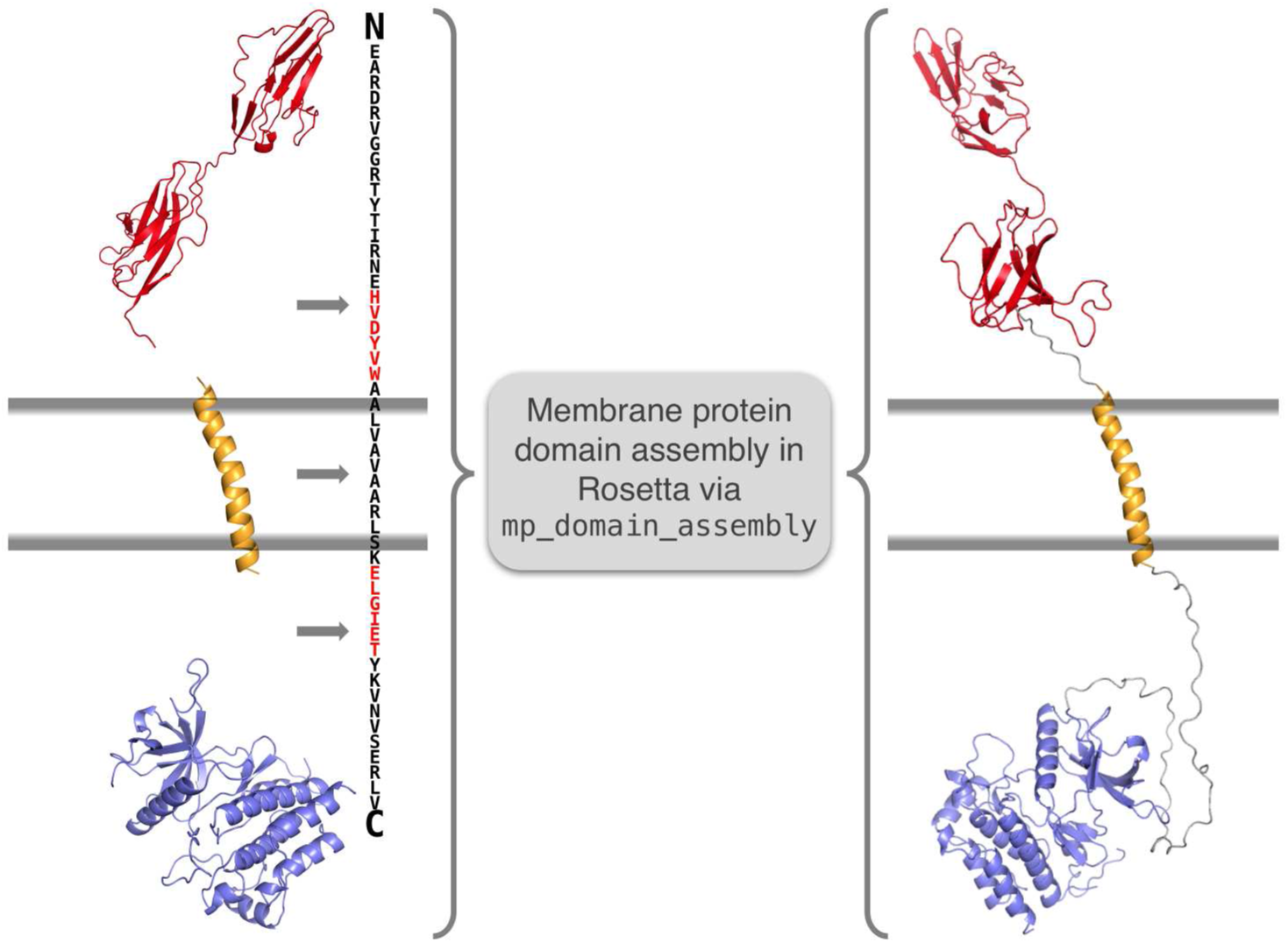
For Table of Contents Use Only:

